# Exploring Correlation-Based Brain Networks with Adaptive Signed Random Walks

**DOI:** 10.1101/2023.04.27.538574

**Authors:** Roberto C. Sotero, Jose M. Sanchez-Bornot

## Abstract

The human brain is a highly connected network with complex patterns of correlated and anticorrelated activity. Analyzing functional connectivity matrices derived from neuroimaging data can provide insights into the organization of brain networks and their association with cognitive processes or disorders. Common approaches, such as thresholding or binarization, often disregard negative connections, which may result in the loss of critical information. This study introduces an adaptive signed random walk (ASRW) model for analyzing correlation-based brain networks that incorporates both positive and negative connections. The model calculates transition probabilities between brain regions as a function of their activities and connection strengths, dynamically updating probabilities based on the differences in node activity and connection strengths at each time step. Results show that the classical random walk approach, which only considers the absolute value of connections, underestimates the mean first passage time (MFPT) compared to the proposed ASRW model. Our model captures a wide range of interactions and dynamics within the network, providing a more comprehensive understanding of its structure and function. This study suggests that considering both positive and negative connections, has the potential to offer valuable insights into the interregional coordination underlying various cognitive processes and behaviors.

## Introduction

The human brain is an intricate and highly connected network, with numerous regions displaying complex patterns of correlated and anticorrelated activity (Bressler and Menon 2010). Functional connectivity matrices (Friston 1994) represent the statistical relationships between these activity signals from different brain regions, often derived from neuroimaging data such as functional Magnetic Resonance Imaging (fMRI), Electroencephalography (EEG), or Magnetoencephalography (MEG). These matrices can be used to investigate the organization of brain networks and to identify patterns of connectivity associated with different cognitive processes or brain disorders by means of various analytical techniques, such as graph theory and complex network analysis (Bassett and Sporns 2017; Bullmore and Sporns 2009).

When performing complex network analysis on correlation-based functional connectivity matrices, one of the following approaches is typically used: 1) Binarization: this involves binarizing the functional connectivity matrix by assigning “1” to connections with correlations above a specified threshold and “0” to connections below the threshold (Bassett and Sporns 2017; Kazeminejad and Sotero 2019). This approach also discards the negative correlations and can be used for further network analysis on a binary network. 2) Thresholding: a more general approach than where we apply a threshold to the functional connectivity matrix, preserving only the connections with correlations above a certain value (often positive correlations) (van den Heuvel and Hulshoff Pol 2010; Kazeminejad and Sotero 2019). This way, we create a binary or weighted network, discarding the negative correlations. This approach can lead to a loss of information about the negative interactions between brain regions (Rubinov and Sporns 2011). 3) Separate analysis of positive and negative connections. Two networks are created – one for positive correlations and another for negative correlations – and then perform network analysis on each network independently. This approach preserves the information about negative interactions between brain regions but treats them as separate entities (Sotero et al. 2020). 4) Disregard information about the connection sign by taking the absolute value of the connections (Meszlényi et al. 2017; Mahadevan et al. 2021). By disregarding the sign of the connections, we may inadvertently overlook critical aspects of the brain’s organization, such as the balance between correlated and anticorrelated activity, which can provide insight into the interregional coordination underlying various cognitive processes and behaviors.

One approach that has shown promise in exploring the structure of complex networks, is the use of random walks (Noh and Rieger 2004). Random walks represent a type of stochastic process that characterizes the random motion of an object or particle in either discrete or continuous spaces (Pearson 1905). Within graph theory and network analysis, random walks are frequently employed to model the movement of a “walker” on a graph, with the walker traversing from one node to another along the edges of the graph (Riascos and Mateos 2021).

Random walks have been adapted for use with signed social networks by adjusting the transition probabilities to preferentially favor positive edges over negative edges, or vice versa (Wan et al. 2019; Zhou et al. 2018; Yang, Cheung, and Liu 2007). However, the interpretation of signed connections in social networks, in terms of “trust” and “distrust”, as well as the associated transition probability models, are not directly transferable to correlation-based brain networks.

As such, a signed random walk model that aligns with the physiological interpretation of correlated and anticorrelated activity in the brain is necessary.

In this study, we introduce a novel adaptive signed random walk (ASRW) approach to analyze correlation-based brain networks, investigating the transition probabilities between nodes (brain regions) as a function of their activities and connection strengths. The distinguishing feature of this model is the inclusion of both positive and negative connections when calculating transition probabilities. Positive connections represent a higher probability of transitions between nodes exhibiting similar activities, while negative connections correspond to a higher probability of transitions between nodes with dissimilar activities. By dynamically updating the transition probabilities in response to node activity differences at each time step and considering connection strengths, the ASRW model offers a more comprehensive insight into the network’s characteristics. To quantify this, we compute the mean first passage time (MFPT).

MFPT measures the average number of steps it takes for a random walker to travel from one node to another in a network. It helps in understanding the accessibility and navigability of the network. Our findings indicate that the classical random walk approach, which considers the absolute value of connections produces lower MFPT than our ASRW model, which considers both positive and negative connections. The proposed method captures a wide range of interactions and dynamics within the network, providing a more comprehensive understanding of its structure and function.

## Materials & Methods

### Computing the functional connectivity matrices

In this study, resting-state (rs)-fMRI data from 491 neurotypical subjects were obtained from the ABIDE Preprocessed Initiative (Cameron, Yassine, et al. 2013). To ensure that our results were not influenced by any custom preprocessing pipeline, we utilized the preprocessed data provided by ABIDE in the C-PAC pipeline (Cameron, Sharad, et al. 2013). The preprocessing involved several steps. First, the skull was removed from images using the Analysis of Functional Neuro Images (AFNI) software (Cox 1996). The brain was then segmented into three tissues using FMRIB Software Library (FSL) (Smith et al. 2004) and the images were normalized to the MNI 152 stereotactic space (Grabner et al. 2006; Mazziotta et al. 2001) using Advanced Normalization Tools (ANTs) (Avants et al. 2011). Functional preprocessing included motion and slice-timing correction, as well as voxel intensity normalization. Nuisance signal regression was also performed, which encompassed 24 parameters for head motion, CompCor (Behzadi et al. 2007) with five principal components for tissue signal in cerebrospinal fluid and white matter, and linear and quadratic trends for low-frequency drifts. Global signal regression (GSR) was applied to remove the global mean from signals, and images were co-registered with their anatomical counterparts using FSL. Subsequently, the images were normalized to the MNI 152 space using ANTs.

The average voxel activity in each Region of Interest (ROI) of the anatomical automatic labelling (AAL) atlas (Tzourio-Mazoyer et al. 2002) was extracted as the time series for that region. As rs-fMRI primarily focuses on low-frequency fluctuations (< 0.1 Hz), a bandpass filter was applied to frequencies between 0.01 Hz and 0.1 Hz. To construct the brain network, the time series for each atlas ROI were correlated with the other regions using Pearson correlation. The strengths of these correlations were used as the connection strengths between different ROIs. In graph terms, this represents a graph where each node is at the center of the corresponding ROI, and each edge weight corresponds to the correlation between the two nodes at opposite ends of the edge.

### Walk rule on the correlation network: the ASRW model

First, let us consider a weighted network consisting of *N* nodes. We place a large number (*K*) of random walkers onto this network, where *K* is significantly larger than *N*. At each time step, the walkers move randomly between the nodes that are directly linked to each other, with transition probability only depending on the absolute value of the connections:

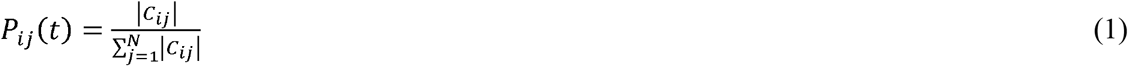

We allow the walkers to perform *T* time steps, and as a walker visits a node, we record the number of walkers in the node. After *T* time steps, we expect that nodes with high degree (hubs) will have a higher number of walkers than nodes with low degree.

For correlation networks, such as the functional connectivity matrix representing the brain network, we propose a routing strategy that takes into account the sign of the connection. In the proposed ASRW model for correlation networks, the transition probabilities are computed based on the weights in the functional connectivity matrix (*C*) and a similarity measure between the activities of the source and destination nodes. The main idea is to favor transitions between nodes with similar activities while also considering the positive or negative weights in the network.

We define the activity in each node *A*_*i*_ (*t*) as the number of walkers visiting the node, and the walkers’ connectivity matrix *W*(*t*), where *W*_*ij*_(*t*) is the total number of walkers that traveled from node *i* to node *j* until time step *t* .

At the beginning (*t*= 0), we use a uniform random distribution to assign each of the *K* walkers to one of the 116 brain areas, resulting in an initial activity *A*_*i*_(0) for each node *i*. Given the activities *A*_*i*_(0) the transition probabilities *P*_*ij*_(t+ 1) and the next activities *A*_*i*_ (*t*+ 1) are computed as follows:

1. For each node *i*, we calculate the adjusted weights for all its neighboring nodes *j* as follows:

a. We introduce the similarity measure *S*_*ij*_(*t*) between the activity of node *i* and node *j* at time *t* using the formula:

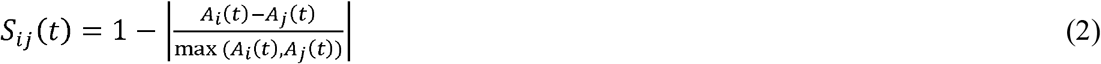

To normalize the difference between the activities of the nodes, we divide by the maximum of the activities max (*A*_*i*_ (*t*),*A*_*j*_(*t*)). The similarity *S*_*ij*_ (*t*) ranges between 0 and 1. A value closer to 1 indicates that the activities of nodes *i* and node *j* more similar, while a value closer to 0 indicates that the activities are more dissimilar.

b. If the weight *C*_*ij*_ is positive, the adjusted weight is 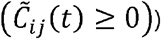:

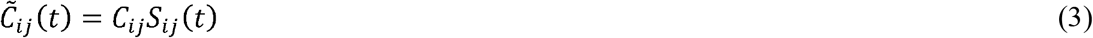

If the weight *C*_*ij*_ is negative, the adjusted weight is:

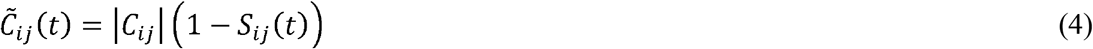

3. We then calculate the sum of the adjusted weights for all neighbors of node *i*. The transition probability *P*_*ij*_(*t*+ 1) of a walker moving from node *i* to each neighboring node *j* in the next time step *t*+ 1 is computed as:

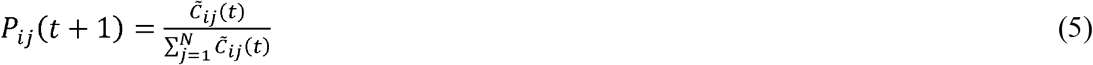

Thus, a walker is more likely to travel through a positive connection *C*_*ij*_ if the activities in nodes *i* and *j* are similar, while it is more likely to take a negative connection if the activities are dissimilar.

4. The random walkers move based on the probabilities *P*_*ij*_(*t*+ 1), and the next activites *A*_*i*_ (*t*+ 1) are the new number of walkers at each node.

5. The walkers’ connectivity matrix *W*_*ij*_(*t*+ 1) is updated by adding the number of walkers that travelled from node *i* to node *j* at time step *t*+ 1 to *W*_*ij*_(*t*), for all pairs of nodes *i* and *j*. Note that although the correlation matrix is symmetric, the resulting walkers’ matrix *W* is usually not symmetric due to the random movement of the walkers.

### Mean First Passage Time Computation

MFPT measures the average number of steps a random walker needs to reach a target node from a starting node for the first time (Hughes 1995). For a network with *N* nodes, the MFPT is the average value of first passage times (*FPT*_*ij*_) over all distinct node pairs, computed as:

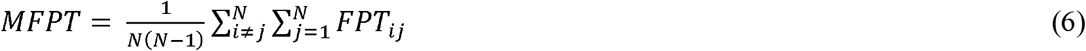

where *FPT*_*ij*_ is the number of time steps walker takes to travel from node *i* to node *j*. To evaluate this, we employ a Monte Carlo approach. Walkers are initially positioned at random nodes and are moved according to a probability distribution defined by the network’s connection weights - this is determined by equations (1)-(4) for an adaptive signed random walk and equation (5) for a classic random walk. Each time a walker reaches a new node for the first time, we record this event’s time. After a defined number of time steps, the recorded first-passage times are used to compute the average, excluding any pairs where a first passage did not occur, giving us the MFPT. This Monte Carlo simulation is repeated multiple times to provide an average estimate of the time necessary for information to spread through the network.

## Results

For each of the 491 correlation-based functional connectivity matrices, we utilize *K*= 5 * 10^4^ random walkers and allocate them throughout the matrix. We employ a uniform random distribution to assign each random walker to one of the 116 brain regions. Subsequently, the random walkers are allowed to move for a duration of *T* = 1000 steps. At each time step, the walkers moved randomly between directly connected nodes, with the transition probability depending on the number of walkers at each node and the strength and sign of the connections (see equations (1)-(4)). We recorded the number of walkers in each node (i.e., *A*(*i*,*t*)) and the number of walkers that transited through each connection at each time step. This allowed us to compute the walkers’ connectivity matrix *W*, where each connection represents the total number of walkers that transited through that connection during the entire simulation. For comparison, we also ran a simulation with a classical random walk where the transition probability only depended on the absolute value of the connections (see equation 1).

Figure 1 shows the rs-fMRI signals from the 116 brain areas (Figure 1A) and the corresponding correlation-based functional connectivity matrix (Figure 1B) calculated from them for one subject, the corresponding walkers’ activity on each node (*A*_*i*_ (*t*)) in the network when using the absolute value of the connections (Figure 1C, equation (1)), and for the signed case (Figure 1D equations (2)-(5)). We can see from the figure that for the signed case, the activities exhibit more variability than for the ‘Abs’ case. This suggests that considering signed connections may provide a more nuanced understanding of the network dynamics.

**Figure 1.**
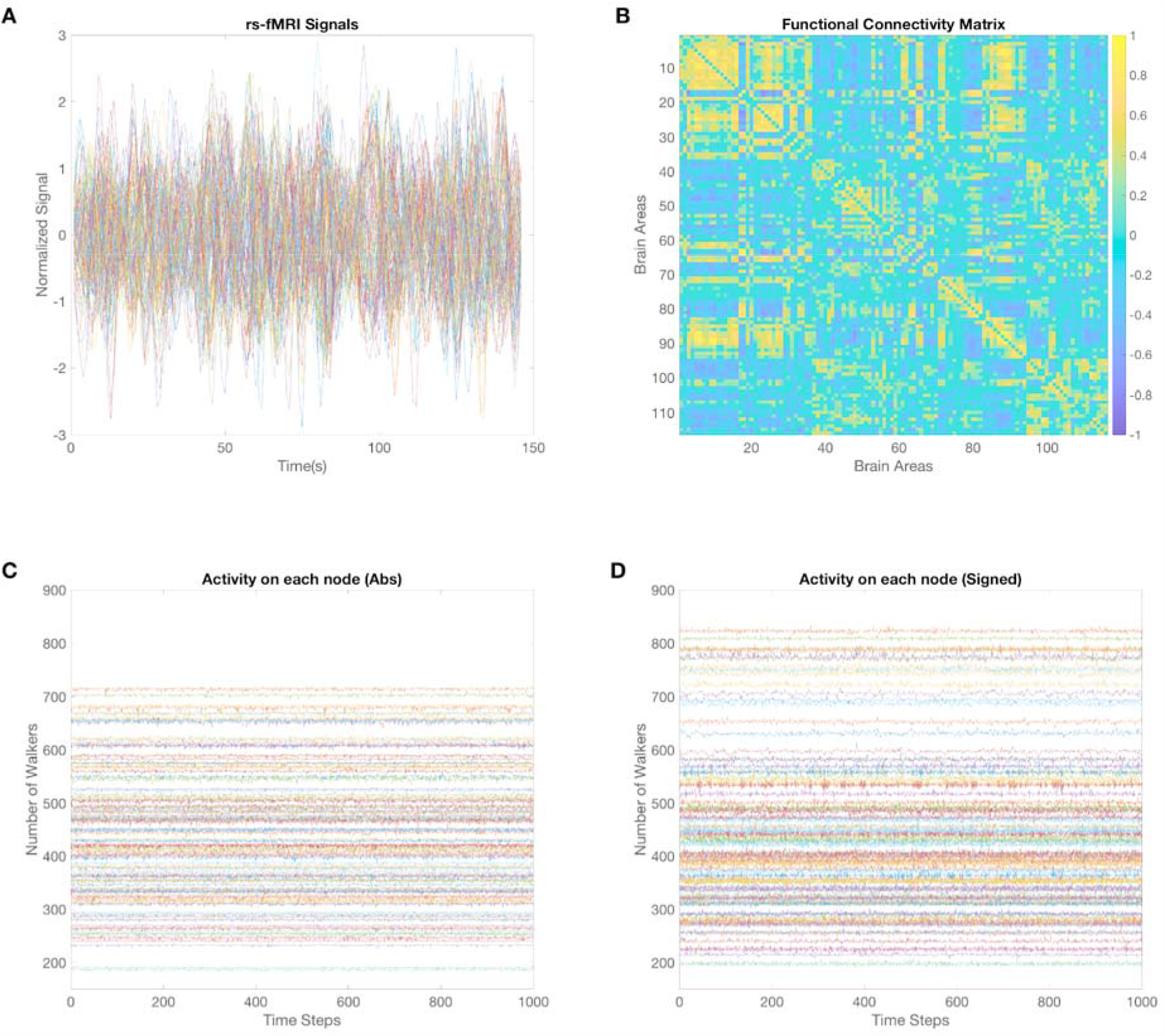
Correlation-based functional connectivity matrix and walkers’ activity on each node in the network for one subject. (A). rs-fMRI signals from the 116 areas in the AAL atlas. (B) Functional connectivity matrix illustrating the strength of correlations between different brain regions for a single subject. (C) Walker activity (A(i,t)) on each node in the network when using the absolute value of the connections, demonstrating relatively uniform activity across nodes. (D) Walker activity (A(i,t)) on each node in the network for the signed case, displaying increased variability in node activity compared to the ‘Abs’ case.

Figure 2 displays the walkers’ connectivity matrix *W*, averaged across subjects for both the ‘Abs’ case (Figure 2A) and the signed case (Figure 2B). We also plotted histograms of the connections in W for the ‘Abs’ (Figure 2C) and signed cases (Figure 2D). In the ‘Abs’ case, we observed 13,334 connections with fewer than 15000 walkers, and only 6 connections with more than 15000 walkers. On the other hand, the signed case presented 272 connections transited by more than 15000 walkers. The signed case revealed a greater number of highly transited connections and demonstrated the presence of highly transited connections between homologous regions in the left and right hemispheres. Specifically, the ten highly transited connections, in descending order, were: 1) ‘Calcarine R’ to ‘Calcarine L’, 2) ‘Calcarine L’ to ‘Calcarine R’, 3) ‘Cuneus R’ to ‘Cuneus L’, 4) ‘Cuneus L’ to ‘Cuneus R’, 5) ‘Cerebellum 7b L’ to ‘Cerebellum 8 L’, 6) ‘Cerebellum 8 L’ to ‘Cerebellum 7b L’, 7) ‘Temporal Sup L’ to ‘Temporal Sup R’, and 8) ‘Temporal Sup R’ to ‘Temporal Sup L’, 9) ‘Temporal Pole Mid L’ to ‘Temporal Pole Mid R’, 10) ‘Temporal Pole Mid R’ to ‘Temporal Pole Mid L’. Here, ‘L’ denotes the left brain hemisphere, and ‘R’ indicates the right hemisphere.

**Figure 2:**
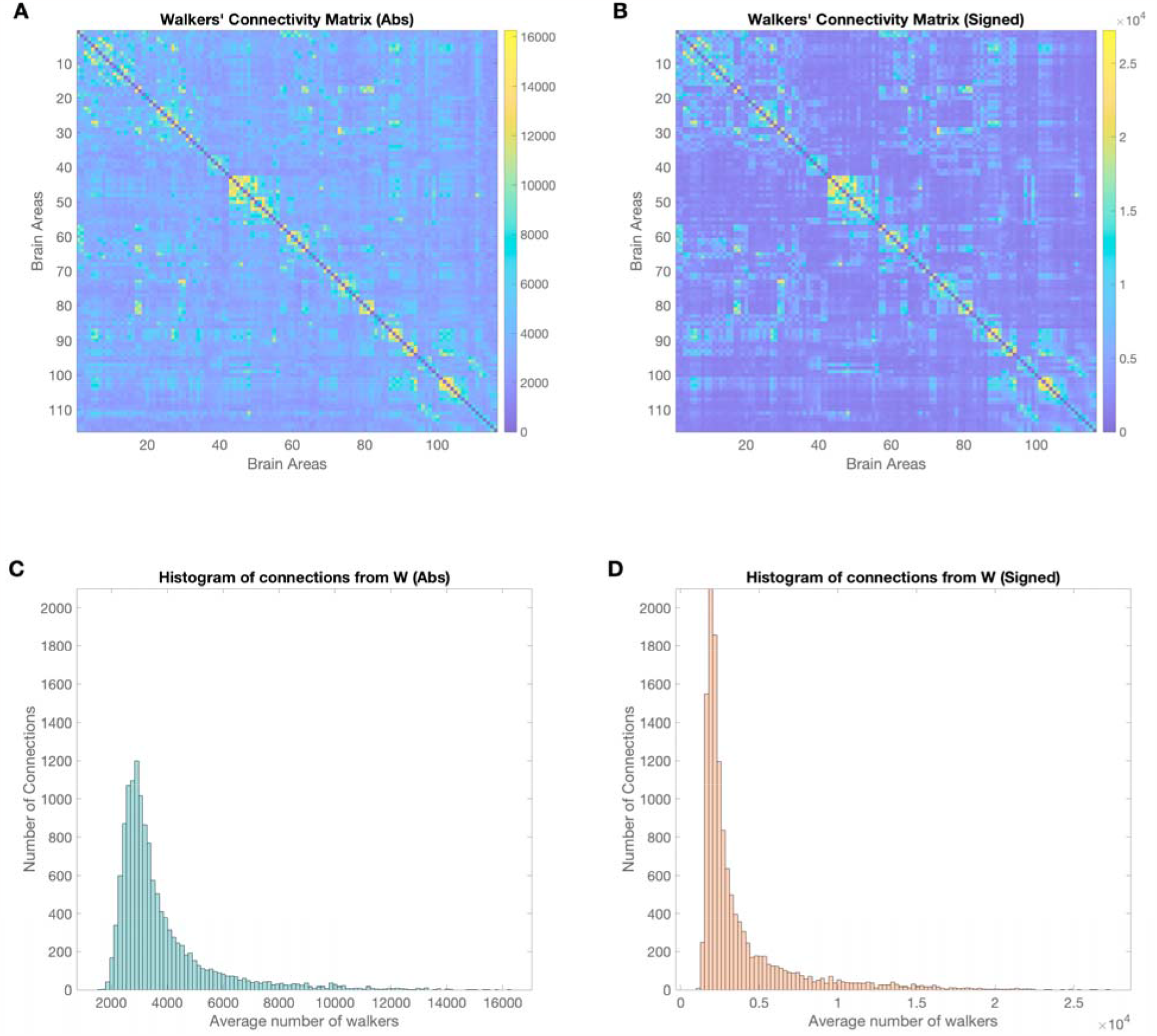
Comparison of walkers’ connectivity matrices and connection histograms for the ‘Abs’ and signed cases. (A) Walkers’ connectivity matrix W, averaged across subjects for the ‘Abs’ case, illustrating the number of walkers transiting between different brain regions using the absolute value of the connections. (B) Walkers’ connectivity matrix W, averaged across subjects for the signed case. (C) Histogram of connections in W for the ‘Abs’ case, showing the distribution of connections with different numbers of walkers. (D) Histogram of connections in W for the signed case, with a greater number of highly transited connections than the ‘Abs’ case.

Figure 3 shows the total number of walkers that transited each brain area, averaged across all subjects and its standard deviation. In the ‘Abs’ case (Figure 3A), the most transited brain areas were ‘Temp Inf L’, ‘Lingual R’, and ‘Temporal pole Mid L’. Interestingly, for the signed network (Figure 3B), the most transited area was also ‘Temp Inf L’, followed by ‘Temporal pole Mid L’. ‘Lingual R’, which ranked as the second most transited area in the ‘Abs’ network, dropped to the 18th position in the signed case. This result highlights the differences in the most transited brain areas between the ‘Abs’ and signed cases, demonstrating the impact of considering both positive and negative connections on the analysis.

**Figure 3:**
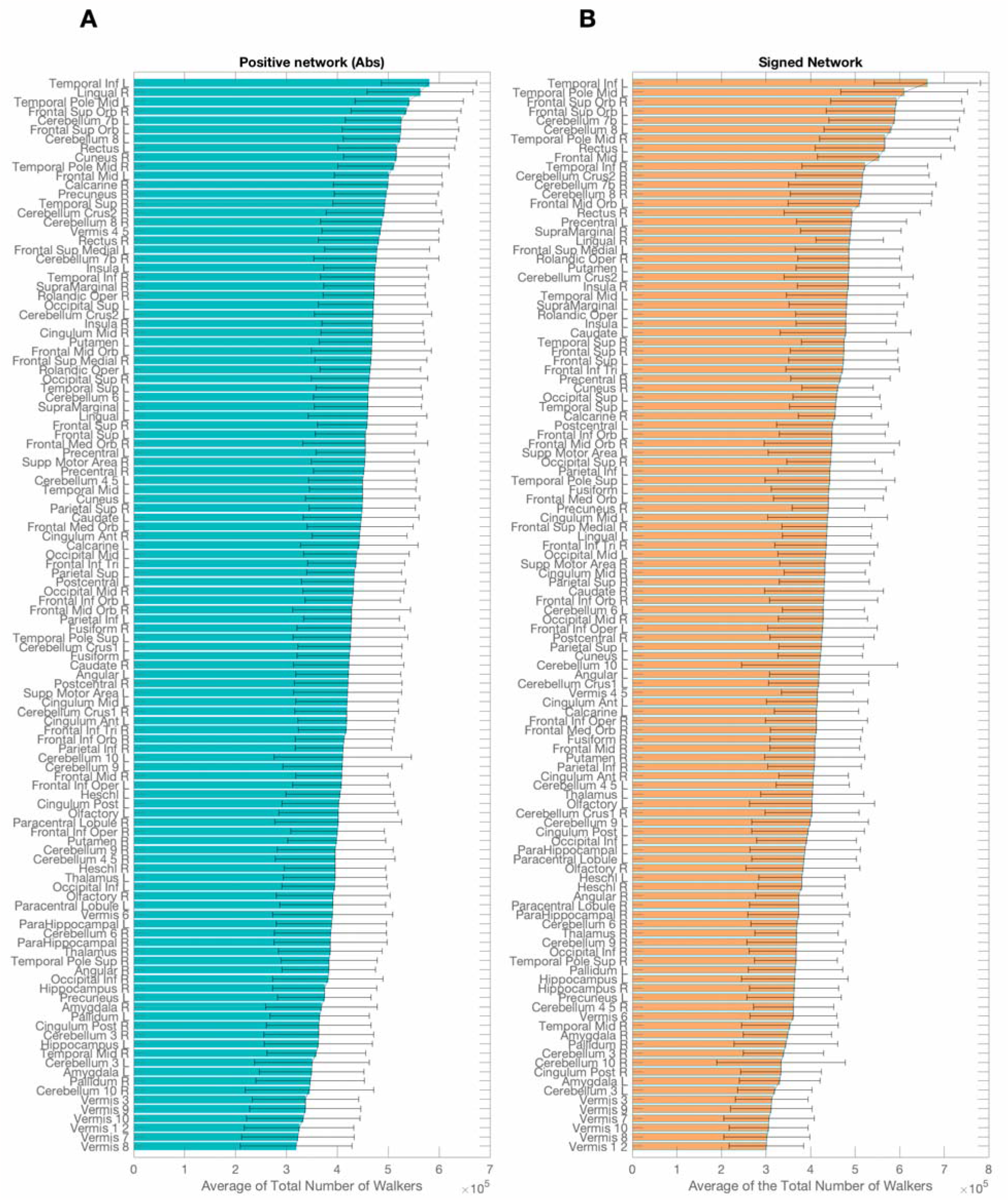
Total number of walkers transiting each brain area, averaged across all subjects for the ‘Abs’ and signed cases. (A) Bar plot showing the mean and standard deviation of walkers transiting each brain area in the ‘Abs’ case. (B) Bar plot illustrating the mean and standard deviation of walkers transiting each brain area in the signed case.

Figure 4A compares the MFPT between the ‘Abs’ and signed cases averaged across subjects. It demonstrates that using the absolute value of the connections results in lower values of the MFPT than the signed case. An important aspect of the signed case is the influence of negative connections in the FC matrix on the MFPT. Figure 4B depicts the MFPT plotted against the percentage of negative connections in the FC matrix for all subjects. It can be observed from the x-axis that the range of negative connections varied from 19% to 40%. We found a significant negative correlation for the ‘Abs’ case (*r* = 0.2728,*pvatue* = 7.87 * 10^−l0^), and nearly identical results were observed for the signed case (*r* = 0.2787,*pvatue* = 3.27 * 10^−l0^).

**Figure 4:**
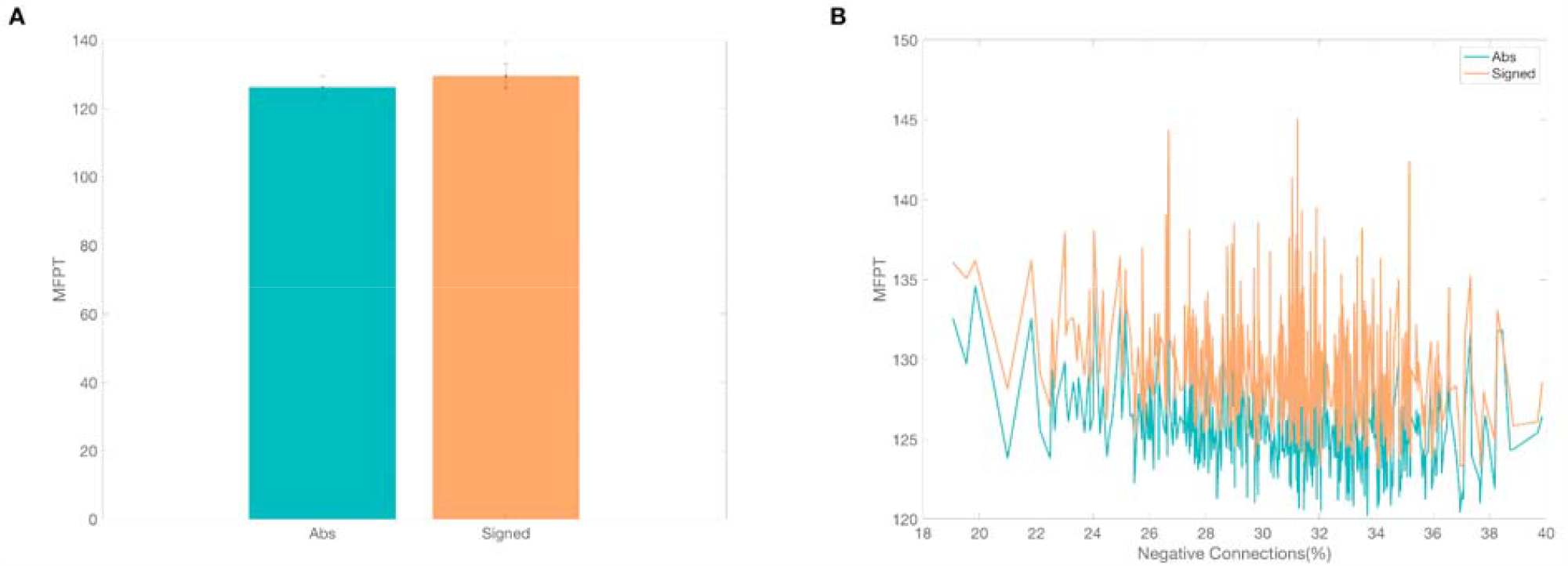
Comparison of MFPT and influence of negative connections. (A) MFPT for both absolute (‘Abs’) and signed connection cases, averaged across subjects. (B) MFPT as a function of the percentage of negative connections in the FC matrix across all subjects.

To interpret these findings, we conducted multiple simulations to calculate the MFPT in a network under different scenarios. We generated a symmetric random network of 100 nodes by creating a symmetric binary adjacency matrix with a designated connection probability (*p*= 0.2)and randomly allocating weights to these connections, while ensuring network symmetry was maintained. We then modulated a variable ‘q’ that determines the fraction of negative connections, ranging from 0 (representing ‘Abs’ or absolute case with no negative connections) to 100%, with increments of 5%. For each ‘q’ value, a signed variant of the random matrix was constructed by selecting and reversing the sign of a fraction ‘q’ of the connections (preserving symmetry to simulate a functional connectivity matrix). Following this, we calculated the MFPT for the resulting signed matrix. This calculation was repeated 200 times for each ‘q’ value, with results averaged. We repeated this entire process for six different numbers of random walkers, denoted as *K*: *K*= 5 * 10^3^, K= 10^4^, *K*= 5 * 10^4^, *K* = 7.5 *, 10^4^, *K*= 10^5^, and *K*= 1.5 * 10S^5^. Figure 5A depicts the outcomes. Our simulations reinforced the empirical data results, indicating that the ‘Abs’ scenario (0% negative connections) yields a lower MFPT compared to cases considering negative connections. Generally, the relationship between MFPT and the percentage of negative connections was non-linear. For relatively low percentages (up to approximately 15%), MFPT increases in line with the percentage. Then, in the range of 15% to 80%, MFPT experiences a slight decrease as the percentage of negative connections increases, which corroborates our findings with real data depicted in Figure 4B. For exceedingly high percentages of negative connections (80%-100%), the MFPT augments along with the percentage increase. Notably, beyond *K*= 5 * 104, subsequent increases in K had a minimal impact on the results.

**Figure 5:**
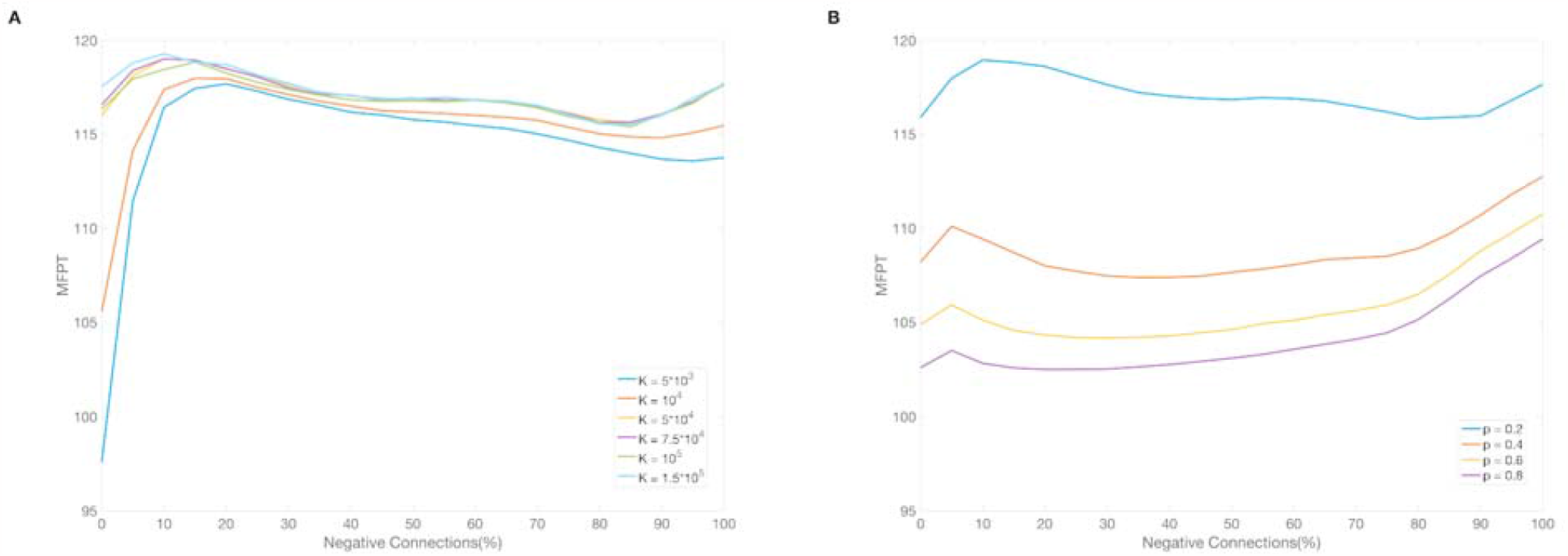
Simulated influence of negative connections, number of walkers, and connection density on MFPT. (A) Influence of negative connections on MFPT for different values of the number of walkers *K*. (B) Influence of negative connections on MFPT for different values of the connection probability *p*.

In a subsequent analysis (Figure 5B), we changed the connection probability, considering four different values: *p* = [0.2,0.4,0.6,0.8], while keeping the number of walkers fixed at *K*= 5* 10^4^. Higher ‘p’ values correspond to denser matrices. Similar to the procedure in Figure 5A, we varied the fraction ‘q’ of negative connections from 0 to 100%, with 5% increments. Each combination of ‘p’ and ‘q’ values (i.e., each simulation) was replicated 200 times, with the outcomes being averaged. Our findings suggest that as ‘p’ increases (hence, increasing the density of connections within the random signed matrix), the MFPT decreases.

## Discussion

In this study, we introduced an ASRW approach for analyzing correlation-based brain networks, taking into account both positive and negative connections. The proposed method captures a broader range of network interactions and dynamics, revealing patterns that might be missed when employing traditional methods. Our results indicate that the classical random walk approach, which relies on the absolute value of connections, underestimates the MFPT compared to the proposed adaptive random walk model. This has important implications for understanding the interregional coordination underlying various cognitive processes and behaviors (Bassett and Sporns 2017).

Despite the advantages of the ASRW model, there are some limitations that should be addressed in future research. One limitation is the reliance on static functional connectivity matrices, which may not fully capture the dynamic nature of brain activity. Future studies could extend the model to account for dynamic functional connectivity, allowing for the investigation of time-varying network organization (Preti, Bolton, and Van De Ville 2017).

An additional aspect to consider in our study is the use of GSR in preprocessing the neuroimaging data. GSR is a common preprocessing step that aims to remove global confounds, such as physiological noise, from the data (Murphy and Fox 2017). However, it has been shown that GSR can introduce more negative correlations into the functional connectivity matrices (Murphy et al. 2009). This has led to some debate regarding the validity of negative correlations as they may, at least partially, be an artifact of the preprocessing technique (Saad et al. 2012).

The potential impact of GSR on our adaptive random walk model is an important consideration, as it could affect the observed transition probabilities and the subsequent network measures. In our analysis, we treated both positive and negative correlations as meaningful connections, reflecting the underlying physiological mechanisms. However, if GSR artificially inflates the number of negative correlations, it could bias our results, potentially leading to an overestimation of the impact of negative connections on the network organization. To address this issue, future research could explore alternative preprocessing techniques that do not introduce additional negative correlations or investigate the sensitivity of the adaptive random walk model to the presence of artificially induced negative correlations. Comparing the results obtained using different preprocessing methods or incorporating additional steps to correct for the possible effects of GSR could help to disentangle the true physiological significance of the negative correlations from the potential artifacts introduced by GSR. This would contribute to a more robust understanding of the brain network organization and its underlying functional interactions.

Finally, future research could also further refine the adaptive random walk model by incorporating additional factors that may influence the transition probabilities, such as the spatial distance between brain regions or the directionality of connections.

## Conclusions

The adaptive signed random walk model presented in this study offers a promising approach for the analysis of correlation-based brain networks, accounting for both positive and negative connections. While there are limitations to the current model, addressing these in future research will help to further enhance its applicability and utility. This method provides a more comprehensive understanding of brain network organization and has the potential to advance our knowledge of the complex interplay between brain regions in various cognitive processes and behaviors.

